# Exploring sources of uncertainty in the estimate of waterfowl harvest in the United Kingdom

**DOI:** 10.64898/2026.05.13.724812

**Authors:** Matthew B. Ellis, Hannah M. Lewis, Tom C. Cameron

## Abstract

There is an urgent need to gather data on harvest rates of waterbirds in Europe to assess the sustainability of hunting. Estimates of total waterbird harvest in the United Kingdom (UK) and the relative harvest of different huntable species come from two separate surveys, the Value of Shooting (PACEC 2014) and National Gamebag Census (NGC, Aebischer 2019), and these have been recently used to explore the likelihood of unsustainable harvests of wild waterbirds by UK hunters (Ellis and Cameron 2022; Madden *et al*., 2025). The reliability of these sustainability estimates depends on how representative the original surveys are of hunter behaviour and success. There are also 1-3 million released game-farm mallard (*Anas platyrhynchos*) that takes up considerable and unquantified proportions of the UK waterbird harvest. Here we explore uncertainties in the UK winter harvest of wild waterfowl by comparing estimates from the NGC dataset with those from the Crown Estate coastal hunting clubs, and a novel approach using analysis of social-media images (2019/20 to 2023/24). We explore the difference in species-specific harvest with and without the uncertainties in the number of released mallard and the total number of duck harvested in the UK. Waterbird harvest estimates differ markedly depending on the input dataset and whether released mallard are included in the analysis. Confidence intervals of each estimate are inflated by uncertainties in the number of released game-farm mallard contributing to, and the size of that national bag. Estimates extrapolated from social media suggest the national harvest of several species may be considerably larger than the corresponding NGC estimates (e.g. Teal *2.07 and gadwall *11.2), while mallard harvests away from formal shoots represented by NGC are significantly lower (*0.71). Excluding released mallard reduces the statistical estimate of total wild duck harvest by 56-63%, which would have biologically significant effects if realised.

## Introduction

There is a need for greater information on the harvest of wild ducks across Europe in order to assess the sustainability of current harvest levels and to inform future management and policy decisions that would attempt to modify the level of harvest (Elmberg *et al*. 2006; Holopainen *et al*. 2018; Aubry *et al*. 2020a). There are two approaches to empirically assess the role of legal hunting on wildlife populations. Mark-resight-recapture approaches can be employed to estimate the survival of individuals, and whether these survival rates would lead to population decline, stability or increase (Folliot *et al*. 2020). These approaches can allow the relative role of hunter vs natural mortality to be assessed (Roberts *et al*. 2023). This approach requires investment in wildlife tagging (e.g. bird ringing/banding) and recording of tagged birds that are resighted or shot. A second approach is to model and predict the harvest rates of different hunted species from the responses to surveys of participating hunters, a practice well developed in North America (Raftovich *et al*. 2025) and in some European nations (Aubry *et al*. 2020a; Lindström *et al*. 2024). Despite the long history of calls for such data (Nicholson 1964; Elmberg *et al*. 2006), there has been remarkably little progress across Europe in developing a standardised approach to collecting hunter harvest data. In Europe there has been some recent development of mobile based applications for hunter reporting, but few states have gone beyond voluntary programs known for significant bias (Aubry et al., 2020). Consequently, in countries such as the United Kingdom (UK) where there is no mandatory requirement for hunters to report their harvest, there are significant challenges in estimating annual harvests, but estimates have been produced using statistical resampling approaches from a range of surveys of hunters or landowners (Aebischer 2019). Using such methods it has been estimated that c1.1 million waterbirds were legally harvested in the United Kingdom in 2016 (Aebischer 2019). Hunter surveys are a standard tool to gather such data and historically these have been attempted by NGO groups in the UK (Harradine 1985). However, producing reliable hunter harvest estimates through surveys is technically challenging (Aubry *et al*. 2020b) and the techniques which produce the most accurate estimates of hunter harvest also tend to be the most costly (Lukacs *et al*. 2011).

There are currently two data sets of hunter harvest data in the UK: the National Gamebag Census and the Crown Estate annual hunting returns (Figure 1). The National Gamebag Census (NGC) is a long term voluntary data recording scheme running for over 60 years and owned and administered by the Game and Wildlife Conservation Trust (Aebischer 2019). Volunteers are landowners or shoot managers, and while they vary in scale they are predominately formal shoots and occur inland (Harradine 1985; Aebischer 2019). Participants to the NGC are reporting on a hunting season’s activity on an area of land rather than direct reports from an individual hunter on the varied hunting activities they may undertake on different properties and locations. With a focus on more formal shooting in larger groups, most often associated as occurring on “estates” and with reference given to birds being driven towards a line of hunters we take this NGC dataset to have a stronger relationship with rear and release gamebird and game-farm mallard shooting than wild bird hunting – but see Aebischer (2019) for a fuller explanation.

**Figure 1:**
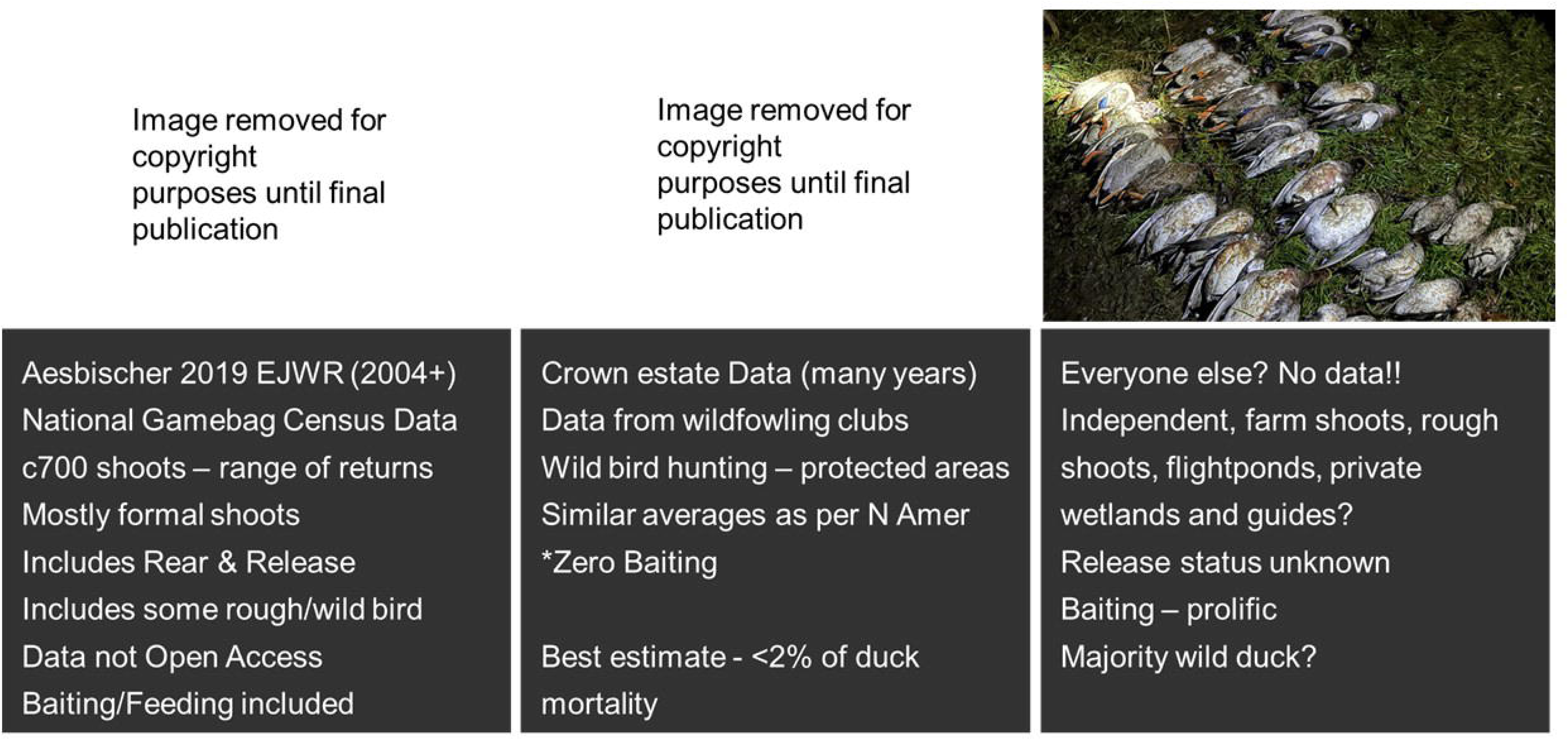
Summary of representativeness of the two existing waterbird harvest data sources as reviewed and determined by the authors. Text summarises sources and for what years or period the data is available, some features of the participants, knowledge gaps and what data management is associated with the source. Note two images removed but will be in final published version.

The Crown Estate is an independent commercial entity responsible for managing land owned by the Monarch of the UK. Crown Estate hunting returns apply when a leaseholder of Crown Estate land, usually coastal wetland, must provide a return to the Crown Estate of the hunting activity each year (Crown Estate 2021). These data are publicly available and mostly apply to what are known as *wildfowling clubs*, almost exclusively this is the hunting of wild waterbirds in more natural landscapes and seascapes, where artificial feeds and baiting is not permitted and hunters are hunting birds exhibiting natural behaviours (Madden *et al.,* 2026). While under scrutiny for most often occurring in coastal protected areas, the contribution of this activity to total waterbird harvests in the United Kingdom is thought to be small (Ellis and Cameron 2022; Madden *et al*. 2025) (Figure 1).

There is a suite of approaches to hunting waterbirds and hunter behaviours that do not fit into either the category of formal duck shooting or wildfowling (Figure 1, left vs. right). These include hunters who hunt waterbirds most often on private lands and unprotected lands such as inland agricultural fields, lakes and flooded pits, floods, wetlands including managed flightponds (“flightponds” are adopted or created wetlands for harvesting waterfowl – where habitat creation, supplementary feeding or baiting, or both are used to attract “wild” waterfowl). In this “unknown” category it is more difficult to assess the role of reared and released game-farm mallards on the bag but it is known to occur at smaller scales on small shoots and private wetlands including flightponds, but it is less prevalent than for supplementing driven shooting days. Releasing game-farm mallard is known to be associated with supplementary feeding or *baiting*, which can attract other waterbirds. Baiting can also refer to the supplementary provision of grain or other feeds in wetlands to attract wild waterfowl to increase harvest success (BASC 2025). It is difficult to assess how widespread baiting is, or its role in affecting harvest of wild waterfowl but based on our own experience and study of social media we believe baiting of wild waterfowl for hunting to be ubiquitous in the United Kingdom. While formal surveys of this third group of independent hunters are not available, the advent of social media and the sharing of hunting images provides a potential alternative to assess relative species-specific waterbird harvests through the use of passive citizen science (Edwards *et al*. 2021).

The use of social media to study social phenomena is well established and there is a growing use of such data in conservation science (Di Minin *et al*. 2015; Levin *et al*. 2015; Lopez *et al*. 2019) leading to the development of more powerful analytical methods (Toivonen *et al*. 2019). Social media has been used to collect socially sensitive data such as on the extent of illegal killing of animals (Eid and Handal 2017) and the extent and structure of the illegal trade of orchids (Hinsley *et al*. 2016), as well as the extent of legal recreational activities such as trophy fishing or if recreation could cause disturbance to wildlife (Lennox *et al*. 2022). Caution is advocated when using such data as social media users may not be representative of the general population (Mellon and Prosser 2017), and there are other well known biases, e.g. prestige bias leading to more pictures or rare or trophy birds. While the most powerful methods of embracing social media tend to rely on users volunteering their information (Aubry *et al*. 2020b; Mneimneh *et al*. 2021), granting additional permissions or interacting in specifically established and monitored fora, this can reduce the pool of users who will choose to interact with such methods and introduce other biases – especially so in more controversial pursuits such as recreational hunting (Hargittai 2018).

Whilst waterfowl harvest data collection is heterogeneous across Europe in terms of effort and methods (ENETwild Consortium et al. 2018; Åhl *et al*. 2021), the paucity of harvest data in the United Kingdom stands in stark contrast to the robust and systematic data collection in countries such as Iceland and Germany. The UK is a major mid-winter wintering ground for many hunted species in Europe, meaning that lack of robust data on the size of the UK harvest of wild waterbirds could be disproportionally more important than for nations where their winter distribution, or estimated harvest is lower. The main quarry species in the UK are the mallard, Eurasian teal (*Anas crecca*), Eurasian wigeon (*Mareca penelope*), gadwall (*Mareca strepera*) and Northern pintail (*Anas acuta*); and three goose species: Canada goose (*Branta canadensis*), greylag goose (*Anser anser*) and pink-footed goose (*Anser brachyrhynchus*). In 2013 there were approximately 28,000 participants in coastal duck and goose shooting (PACEC 2014). Providing similar estimates for inland waterbird shooting is more difficult but expected to be between 75,000 and 280,000 (PACEC 2006). The uncertainty surrounding the number of mallard released further complicates harvest estimates (estimates vary, e.g.870,000 −1.3Million (Aebischer 2024), 0.9 - 6Million (Madden 2021)), especially when there is incomplete knowledge of the population of hunters sampled.

Increasingly we are reliant on accurate and regular wildlife harvest estimates to inform discussions on the sustainability of harvest levels at the national and international levels (e.g. (EU TASKFORCE 2024), and such data will be essential for the proper functioning of adaptive harvest systems such as the African Eurasian Migratory Waterbird Agreement’s (AEWA’s) European Goose Management Platform (https://egmp.aewa.info/) (e.g. Madden *et al*. 2025). To demonstrate the importance of a shift in effort towards a formal data reporting scheme, in this study we assess the role of the currently available United Kingdom harvest data on the estimate of species-specific waterfowl harvests. Specifically, we explore the consequences to estimated UK waterfowl harvest of using the NGC, the Crown estate data or social media derived data as representative of the average UK hunter activity. We do not re-estimate total national waterfowl harvest or test which is the best or more robust method for the species-specific bag data as we have already outlined they all have significant biases, but we use this analysis to demonstrate the uncertainty in these estimates as a further plea for state or voluntary investment in more robust and appropriate methods (e.g. Elmberg *et al*., 2006, Holopainen *et al*., 2018). Finally, we explore the possibility of estimating the demographic breakdown of hunted species statistics, e.g. sex and age, using the same data from photos. Such techniques have the potential to generate low-cost harvest estimates and trends in age/sex structure which could be useful for managers either on their own, or in combination with more formal methods.

## Methods

Digital photographs of hunting bags from two UK specific Facebook groups focused on duck and goose hunting (“Flight Pond Duck Shooting and Conservation” and “Foreshore wildfowling & flight pond Shooting, Management And Conservation”) were analysed for the 2019/20 to 2023/24 hunting seasons. The research proposal to undertake this work using submission to each group as consent to use the data in the photos was approved by the University of Essex ethics committee (ETH2122-0809; see discussion). No human participants were involved in this study. Between January and December 2024 we accessed each Facebook group and viewed and analysed posts remaining on the group for the 2019-2024 period. For each photograph we recorded the date the photograph was submitted, all ducks and geese were attempted to be identified to species and we attempted to age each species using plumage and bill characteristics (Baker 2016; Mouronval 2016). No attempt was made to determine the sex of geese from photographs, but the sexing of ducks was attempted using plumage characteristics as above. Photographs accompanying adverts for hunting opportunities were excluded from the analysis, and where a user submitted multiple photographs on the same day, as part of the same collection of images, only the photograph with the most individual birds was included unless photographs were clearly from different hunting trips (for example different hunters, different clothes or different vehicles). Where users reported a specific number of birds harvested but the accompanying photograph showed fewer individual birds the additional unphotographed birds were recorded as unknown. Where species, age or sex identifications were provided along with the photographs we recorded them and assessed whether they were correct. Finally, where provided, we recorded the number of hunters who contributed to the harvest shown in the photographs and recorded the data as missing if not reported. As a condition of our ethical approval, we did not download and save the photos or retain any information about the identity of the hunter or the user posting the photo, and therefore in statistical terms all photos are considered independent. This is not ideal, likely incorrect and while it may not influence mean harvest estimates it will underestimate confidence intervals.

We inferred the total number of ducks and geese shot in the UK using total harvest estimates from the UK Value of Shooting Report (PACEC 2014) and attributed to the approximate harvest for 2016 as per previous studies (Aebischer 2019). Estimates of the total UK waterbird harvest have since been updated for 2021/22 (1.1 million waterbirds, Cognisense 2024).But here we use the 2016 estimates for total harvested waterbirds as they are split by ducks and geese and the difference in total harvest between 2012, 2016 and 2021 will make no qualitative difference to our results (i.e. 1.1-1.2 million). In summary, 95% confidence limits for the numbers of different species of ducks and geese shot in the UK were obtained by bootstrapping at the level of photos, with 10,000 bootstrap runs. Previous studies treated the group estimates ±10% as 95% confidence limits (Aebischer 2019), and so we obtained confidence intervals for the estimates by dividing at least 10% of the estimate by 1.96 depending on the required distribution. For each bootstrap run we resampled with replacement a number of photographs equal to the original sample and calculated the proportion of each species present in the resample. We applied the proportions of each species to a value drawn from a normal distribution with a mean equal to the published total estimate for “ducks” and “geese”. The 95% confidence limit was taken to be the lower and upper 95^th^ percentile of each of the per-species estimates from the 1,000 bootstrap runs.

The proportion of each species in the bag was estimated from the only two additional contemporary sources. Aebischer’s estimates for 2016 (Aebischer 2019) were based on participants in the National Gamebag Census and as previously discussed are likely to represent hunters engaged in more formal and/or commercial hunting. The Crown Estate wildfowling returns make no attempt to estimate national totals (Crown Estate 2021), but report the annual returns of approximately 150 duck and goose shooting clubs and generally fewer than 1,000 individual duck and goose hunters who hunt on the coast. In the absence of complete harvest data for the UK we are not making an assessment on which of these estimates is closest to the true relative species-specific harvest on UK waterfowl, but are instead exploring the differences in species specific harvest estimates generated from different data sources.

In a second analysis, we considered ducks only to explore further sources of uncertainty for estimating species specific harvests. Estimates of the proportion of the UK mallard harvest attributable to reared and released game-farm mallards were taken from recent analyses of National Gamebag Census (NGC) data (Aebischer 2024). We multiplied estimates of the harvest at release sites as a proportion of the number of mallards released (i.e. return rates) for four years (2002, 2012, 2016 and 2022; range 61.6–64.3%, mean 63.0, variance 1.3) by the total number of mallards released to obtain an estimate of the contribution of released mallards to the national mallard bag in a given year. These return rates are subject to uncertainty. It is likely that some birds harvested on release sites are wild, and harvest efficiency estimates derived from the NGC may contain sampling biases (see Discussion). Likewise used in the way we do here they could result in underestimates of the total number of game-farm mallard in the total duck bag if large numbers of game-farm mallard are shot away from release sites or the estimate of total released ducks used in our analysis is wrong (e.g. Aebischer vs Madden estimates). Therefore, we next explored combining the uncertainty in return rates combined with the uncertainty in total game farm mallard releases (mean 1.2 million, 0.88-1.6 million; Aebischer 2024). Finally, we explore the uncertainty in the total national UK harvest of ducks (mean 1.135 million, 0.83-1.52million ducks, Aebischer 2019). While this last analysis shall not change the mean harvest estimates, it is useful to consider how it changes confidence intervals for comparison to the NGC estimates of Aebischer (2019).

Using the same image-based approach as for all waterbirds, we repeated the analysis for ducks only but incorporate uncertainty into the bootstrap procedure by random resampling from a distribution with the same mean and variance as the NGC return-rate estimates in each bootstrap run (uniform distribution, mean 0.63, s.d 0.013). This resampled release rate was multiplied by the estimated total number of mallards released in 2016 either as a fixed value of 1.2 million ( Aebischer 2024) or by resampling from a distribution with the same mean and variance as the NGC estimates of released game-farm mallard (normal distribution, mean 1.2 million, s.d. 86326) to estimate the number of released mallards harvested in that year. We explored uncertainty in return rates from the NGC being representative by drawing return rates into the bootstrap procedure by resampling from a distribution that represents rates across 6 experimental shoots we work with (normal distribution, mean 0.5, s.d. 0.03 – giving appx. range from 40-60% return rates).

Whichever method, the estimate of total released ducks harvested was then deducted from the total *number* of ducks harvested in 2016 as either a fixed value of 1.135 million or from a distribution with the same mean and variance as sum of lower or upper confidence intervals of duck species in the NGC estimates (excluding Goldeneye, mean 1.135 million, s.d. 193979 – giving an approximate range from 0.88 to 1.515 million ducks Aebischer 2019). The remainder is assumed to represent the total harvest of wild ducks of all species.

Finally for each method, the bootstrapped proportions of each species found in the social media image analysis were applied to the run’s value for the total number of only wild ducks of all species harvested in 2016. For each species, 95% confidence limits were taken to be the lower and upper 95th percentile of the bootstrap distribution from 10,000 runs.

All analyses were conducted in R (v4.5.1) (R Core Team 2025).

## Results

A total of 682 social media photographs were analysed containing 5,407 birds, of which 5% could not be identified, 30% were geese, 0.5% were waders or rails (golden plover, snipe, woodcock and coot) and the remainder ducks. All of the legally shootable duck and goose species were recorded at least once, with the exception of goldeneye (*Bucephala clangula*). The mean number of waterbirds present in the photographs was 8.3 (SE = 0.4) with 14.7% of photographs containing only a single bird (Figure 2). The greatest number of birds present in any photo was 96 mallard. All unidentifiable birds were ducks and typically could not be identified as they were reported but not photographed, or due to poor quality photographs including blurring, poor lighting and piling of birds. Sex determination was possible for 65% of those ducks we could identify to species (Table 1), but age determination was only possible for 3% of ducks and 1% of geese and no further data is presented for this.

**Figure 2:**
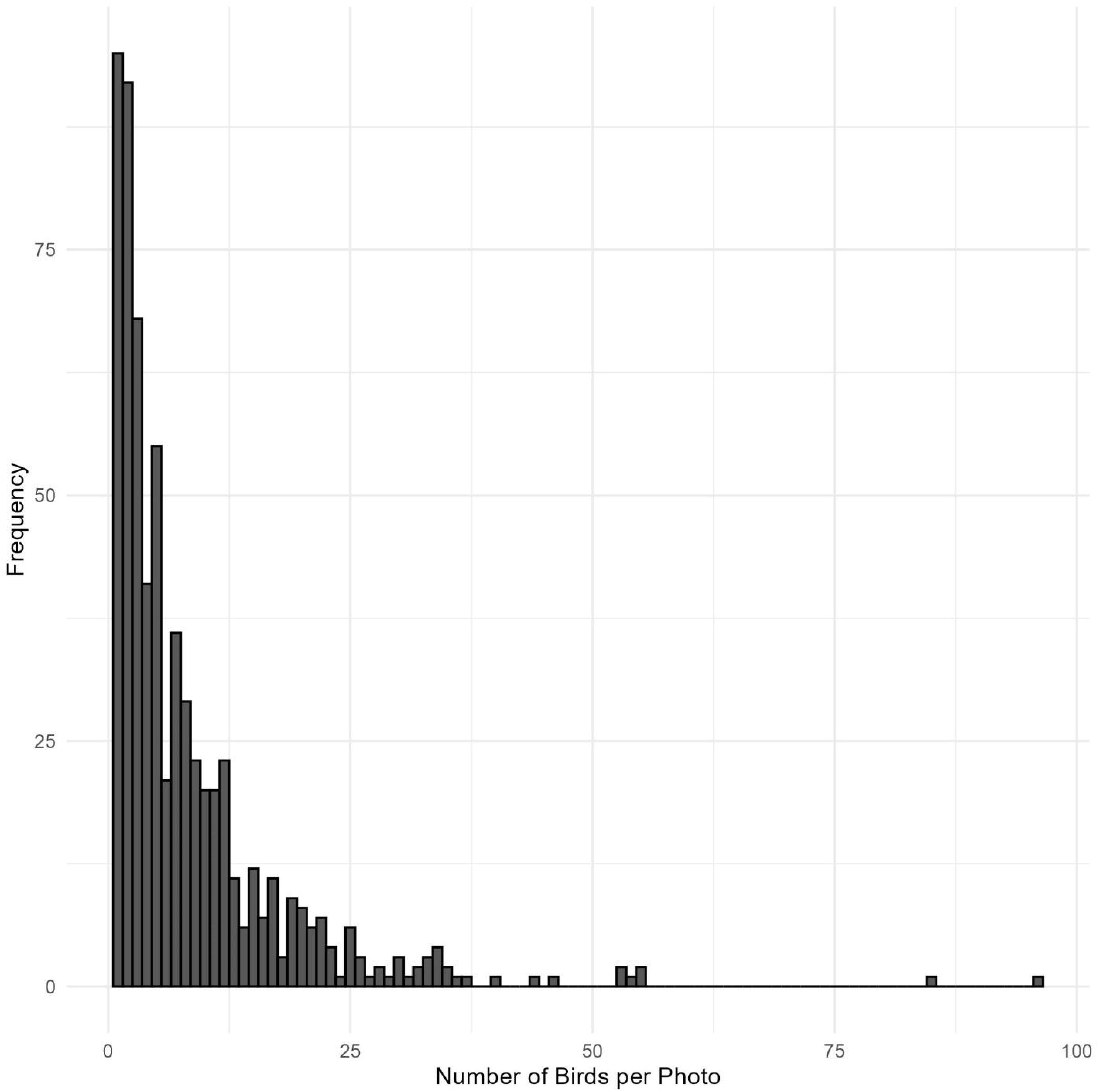
Distribution of waterfowl bag sizes from 271 analysed photographs of waterbird harvests. This includes all images that incorporated solitary and group hunting, commercial inland goose guide hunts and non-commercial do-it-yourself hunting both inland and at the coast.

**Table 1.** The total number of ducks identified, the percentage assignable to a sex class and the percentage of male birds per species.

| Species | n | Percentage |  |
| --- | --- | --- | --- |
|  |  | Sex determined | Male (of those possible to determine) |
| Mallard | 2,041 | 83.9 | 50.8 |
| Teal | 878 | 47.3 | 31.1 |
| Wigeon | 271 | 54.2 | 43.5 |
| Gadwall | 167 | 50.9 | 38.9 |
| Tufted duck | 54 | 13.0 | 12.9 |
| Shoveler | 37 | 78.4 | 48.6 |
| Pintail | 21 | 85.7 | 66.7 |
| Pochard | 8 | 100.0 | 62.5 |

Species identification was provided by the authors of 3.5% of photographs and all identifications were correct. The number of hunters was provided in 51.5% of photographs, and the average number of hunters in each of those was 1.6, with on average 4.5 ducks or geese per hunter.

The percentage of each species in the UK waterfowl bag (split by ducks and geese) from our social media analysis is shown alongside that of NGC and the Crown Estate estimates (Table 2). Social media analysis suggests that proportionally fewer mallard (59 vs. 83%) but more teal (25 vs. 12%), wigeon (7.8 vs. 4%) and gadwall (4.8 vs. <1%) are harvested by UK hunters represented by posts on social media compared to estimates based on the NGC (Table 2). Compared to the Crown Estate input data, our analysis suggests the average UK waterfowl hunter posting on social media bags proportionally more mallard (59 vs 24%), but fewer teal (25 vs. 37%) and fewer wigeon (7.8 vs 34%) than a coastal wildfowler.

**Table 2:** Estimated harvest of duck and goose species in the UK and the estimated percentage of the harvest per group (given separately for the total harvest of ducks (1.135 million) and geese (110,000)) from social media, NGC and Crown Estate input data. Harvest estimates for Goldeneye were not determined as there were no social media images submitted of this species; therefore estimate is effectively 0(0,0). Where confidence intervals in the estimate from our analysis of social media images and those of Aebischer (2019) overlap – they are denoted by an asterix (*). Where the mean estimate from analysis of social media images does not overlap with the confidence intervals in the NGC estimates (Aebischer 2019), these are denoted by ^.

| Species | Estimate (95% CI) | Percentage | Percentage |  |
| --- | --- | --- | --- | --- |
|  |  | Facebook<br>2022-24 | NGC 2016<br>(Aebischer<br>2019) | Crown Estate<br>(2021) |
| Mallard | 670,000 (580,000 – 760,000)*^ | 59 | 83 | 24 |
| Teal | 290,000 (230,00 – 340,000)^ | 25 | 12 | 37 |
| Wigeon | 89,000 (60,000 – 120,000)*^ | 7.8 | 4 | 34 |
| Gadwall | 55,000 (39,000 – 74,000)^ | 4.8 | <1 | 1 |
| Tufted duck | 18,000 (9,000 – 28,000)*^ | 1.5 | <1 | <1 |
| Shoveler | 12,000 (7,600 – 18,000)^ | 1.1 | <1 | <1 |
| Pintail | 6,900 (3,700 – 11,000)^ | <1 | <1 | 3 |
| Pochard | 2,600 (330 – 6,100)*^ | <1 | <1 | <1 |
| Goldeneye | ND | <1 | <1 | <1 |
| Canada goose | 39,000 (29,000 – 50,000)* | 35 | 28 | 73 |
| Pink-footed<br>goose | 36,000 (25,000 – 46,000)* | 33 | 11 | 5 |
| Greylag goose | 35,000 (26,000 – 40,000)* | 31 | 61 | 22 |
| White-fronted<br>goose | 420 (50 – 990)*^ | <1 | <1 | <1 |

The mean estimate of number of the 1.135 million ducks harvested in 2016 that are game-farm released mallard is 756,000, reducing the total estimated harvest of *wild* ducks to 379,000. The relative proportions of each wild duck species harvested in this second simulation based on social media data is the same as the previous model, but when all these proportions are converted to estimates of the national UK harvest large differences emerge in the mean and confidence intervals when sources of uncertainty in game farm mallard releases, return rates and total harvest are explored (examples shown in Figure 3). For teal the estimate can change from 140,000 based on the NGC, to 290,000 using proportions from social media analysis, or as low as 96,000 using social media if we first remove the estimated proportion of released mallard (Figure 3). This same approach means that wigeon harvest estimates vary between 89,000 (social media), 43,000 (NGC) and 30,000 (social media without releases; Figure 3). Similar proportional changes occur for others species – including those of current conservation concern (e.g. pochard, pintail and shoveler) – but their harvest estimates always remains small (see Supplementary Information). Our model simulations with social media and game-farm release estimates suggests 220,000 – 310, 000 wild mallard were harvested in the UK in 2016 (see Table S1 for CIs for each of these estimates).

**Figure 3:**
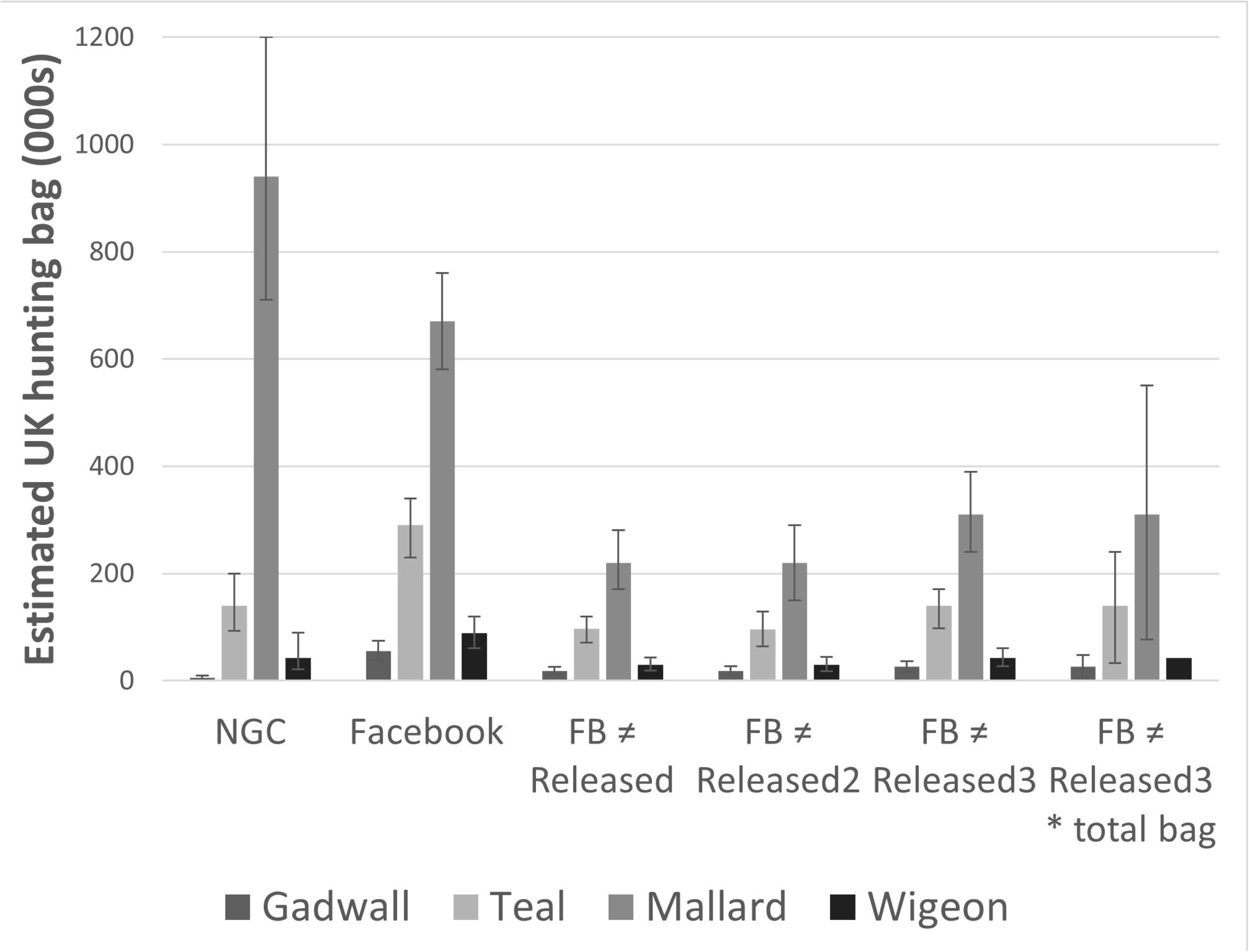
Mean estimate of harvest of four of the most common hunted ducks in the United Kingdom in 2016 repeated from the National Gamebag Census (NGC, Aebischer 2019) that is known to include data from survey respondents that release game-farm mallards, and that same estimate when based on the proportions as seen on social media (Facebook), or the bag when assuming that for ducks those birds seen on social media are not linked to commercial hunting or rear and released game-farm mallards using estimates based on return rates from Aebischer (2024) (FB ≠ Released), or when release estimates are based on uncertainty in return rates from our own work (FB ≠ Released2) or in addition to FB ≠ Released2 including uncertainty in the total number of game-farm mallard released (FB ≠ Released3), or finally in addition to FB ≠ Released3 including uncertainty in the total number of wild duck harvested in the United Kingdom (FB ≠ Released3* total bag). Error bars are 95% confidence intervals.

## Discussion

Estimates of the relative proportion of each species of ducks and geese harvested by hunters in the UK differ markedly depending on the data source (i.e. harvest data) and are heavily influenced by the inclusion of released mallards in the estimates of the total number of waterbirds harvested. The mean proportions of each species in the bag as generated from analysis of social media posts tended to fall between those made from NGC data and the Crown Estate data. However, these differences are largely driven by the proportion of mallard in the bag which varies from 24% on the coast (Crown Estate data) to 83% for the more formal inland shoots associated with releasing game-farm mallard (NGC). Social media based mean bag estimates are predominately higher than the NGC and sit outside the confidence intervals of those from the NGC based estimates for most of the huntable fully wild duck species for which we were able to gain a statistical estimate (Table 2). For mallard, Tufted duck, wigeon and Pochard the confidence intervals of waterbird harvest estimates of the two methods overlap, but not for teal, Gadwall, shoveler and pintail. Other place-based factors are likely to have influenced far higher proportions of teal and wigeon (e.g. 37 and 34% respectively) in the estimates from the Crown Estate data; but this is to be expected given the association between these wild migrant waterfowl and coastal saltmarshes. While these high percentages of wigeon in Crown Estate bags may be alarming, the total numerical contribution of Crown Estate hunting to the national bag is thought to be very low (c<2% Ellis and Cameron 2022).

It is not possible to determine whether the mallard reported to NGC or Crown Estate or present in social media photos are genuinely wild or part of the cohort of c1.2-2.3 million mallard released in the UK each year by shoots, clubs and landowners (Aebischer 2019; Madden 2021; Aebischer 2024). Our expectation is that a large majority of the mallard shot in the UK are reared and released game-farm birds, and they are shot by a minority of UK hunters, or minority of hunter effort, and are most represented by the hunting activities and practices employed by typical respondents to the NGC. However, some of the social media pictures we analysed included large bag sizes of mallard alongside pheasant, with for example the largest bag containing 96 mallard, which are bag sizes much more likely to be associated with reared and released game-farm mallards. It could then be argued that we should include rear and released mallards in any assessment of the sustainability of wild waterfowl hunting in the UK. Unfortunately any inclusion of harvested game-farm mallards inflates the estimates of the total number of harvested waterfowl for all species, i.e. their estimate is a proportion of 1.135 million harvested ducks instead of a proportion of a much smaller number.

As a first step towards determining the role of inflated hunting bag statistics from released mallard we estimated the number of each hunted duck species harvested in the UK with and without an estimate of the released game-farm mallards in the UK hunting bag. This has a huge effect on the statistical estimates of wild duck shot in the UK, with far fewer teal (97,000 vs 140,000) and wigeon (30,000 vs 43,000) in particular when compared to the original estimates from the National Gamebag Census (Aebischer 2019). This exercise provides our first estimate of the number of wild mallard harvested in the UK, such that in 2016 when Aebischer (2019) estimated the total mallard bag including released game-farm mallards, to be 940,000 (710,000 – 1,200,000) we estimate 220,000 (170,000-280,000) of these were wild birds. This is supported by estimates of local bag returns from National Gamebag Census participants who released game-farm mallard, as they reported to harvest an average of c63% of the number of mallards they released which amounted to 756,000 released birds; not dissimilar to the 720,000 we estimated in our simulation model (c940K-220K). While these estimates appear to be in the right ballpark – 940,000 vs 756,000+220,000, our lack of accurate estimates of the proportion of game-farm released mallards in the national bag can clearly have a huge influence on the statistical estimates of wild duck species harvest whether that is to bias estimates down by assuming the NGC is representative (i.e. 63% of released ducks end up in the national bag) or bias them up because total game-farm mallard releases and harvest are being underestimated (e.g. harvest of game-farm mallards off site in both release and subsequent years).

Only a quarter of shoots reporting to the NGC that shoot mallard also release mallard (21-27.2%, (Aebischer 2024)). The remaining three-quarters shoot an unknown ratio of wild and released mallards attracted to those locations via habitat management or baiting. Game-farm mallards could therefore make up an even higher proportion of the national bag than we have estimated here, which would consequently further reduce the statistical estimates for other wild species. Movement of game-farm mallards is considered to be reduced compared to their wild counterparts (as reviewed in Aebischer (2024) but see also Madden *et al.,* 2026), but these small few to tens of km movements can still be ecologically meaningful, create connections with important wild bird aggregations and result in significant harvest away from release sites.

We were able to determine the sex of the majority of ducks in the photos. However, this was influenced by the large number of mallard (84% determined) and for the remaining harvested species our rate of successful sex determination was generally low or based on a small number of birds (most below 50% determined). In many cases our ability to determine sex was largely a function of our ability to identify adult males, rather than to determine males per se. In consequence, we have less confidence in the accuracy of the sex ratio estimates for all species except mallard. We were not successful in estimating the ages of either ducks or geese from the photos in our study. The difficulty in both aging and sexing birds was often due to a combination of poor lighting and birds being photographed on their backs, normally resulting in their wings, heads and tails being obscured. If submitted photographs were to be a bag recording method, simple guidance could result in substantial improvements in the proportions that could be successfully aged and sexed (e.g. as per scheme in Latvia (Stīpniece *et al*. 2025)).

Our motivation for this work is not to dispute the quality of the research or procedures used to gain our current estimates for the UK national game bags or numbers of released game-farm mallard. Indeed, our own research has very much depended on it, e.g. (Ellis and Cameron 2022; Madden *et al*., 2025). However, the raw input data from each of the data recording methods we have explored in this study are imperfect, each have inherent biases (Figure 1) and each introduce uncertainty to any estimate of species-specific hunting bags and therefore also any measure of sustainability of that bag. However it should be noted that uncertainty can be accounted for (McGowan *et al*. 2011), and we now have several relatively consistent methods to determine whether these harvests are sustainable or not. Lack of data and appropriate models are not as much of a barrier to adaptive management of UK recreational waterbird harvest as might have been assumed (Johnson *et al*. 2018; Ellis and Cameron 2022, Madden *et al*., 2025). For over two decades researchers, conservation and hunting organisations have asked Eurasian flyway nations to undertake more appropriate data recording, collaboration and adaptive management of wild bird harvest (Elmberg *et al*., 2006; Holopainen *et al*., 2018); exactly as is now celebrated for the recent collaborative actions on Eurasian turtle dove (Carboneras *et al*. 2024). Instead several nations, including the UK and Ireland, have consistently fallen short of this and instead opt for binary judgement for species-specific hunting management (e.g. Wildlife (Wild Birds) (Open Seasons) (Amendment (S.I. No. 421 of 2023)); and as is proposed to be repeated again in the United Kingdom (e.g. UK DEFRA Consultation on Amending the Wildlife and Countryside Act 1981).

Our methods represent a volunteer sample, as individuals can choose to self-report their bag by posting their photos on publicly available social media pages. We do not know if users of these groups are representative of hunters in the UK in terms of hunting activity, success, location or demographics that could influence bag estimates. This is an important limitation of our data as such volunteer sampling can introduce selection bias (Aubry *et al*. 2020b), especially if there is any correlation between the likelihood of self-selecting individual harvest or the type of waterbird hunting activity undertaken. For example, as we set out in Figure 1 it is less likely that those posting pictures of hunted quarry to the social media groups we have studied are posting pictures from more formal commercial *driven* or released duck shoots. We cannot test this assumption using our data as no information about those posting pictures of hunting outcomes was collected, though attempts to compare hunter characteristics including activity, age and experience, all known to affect harvest (Heberlein *et al*. 2002), could be assessed in a future study.

Following publication, it is possible that the use of social media as an independent source of the structure of waterbird harvest in the UK will change. We did not announce our intention to use these photos in this way to these Facebook group members, nor seek direct or specific written or verbal consent for the use of photos. There are good reasons to assume that should we seek permission that we would receive few positive responses. Furthermore, seeking permission may also affect the behaviour of those posting that could enhance externalities such as post-harvest guilt, prestige bias and cultural barriers – factors that will already affect the results we have presented here - thus making future works less representative of the structure of the UK wild waterbird harvest (Farley *et al*. 2021). This approach to consent has been used recently to evaluate uptake of voluntary and legal measures to switch to non-lead ammunition, not using informed consent from landowners as the authors felt it was likely to create significant bias (Green *et al*. 2023). We would encourage UK hunters to not assume ill intent on our part, as we are conducting these studies to find solutions that best manage real and perceived conflicts between wild bird harvesting and conservation obligations of the United Kingdom, shifting the harvest system to one that can demonstrate sustainability or respond to well evidenced concerns if the harvest is unsustainable.

We believe that our method allowed for the rapid collection and assessment of harvest data from a segment of hunters which have proven difficult to accurately assess; individuals and small groups hunting wetlands, coastal but also inland, noting the vast majority of wild waterbird hunting effort occurs inland (i.e. 28,000 shooting days for coastal hunting vs 38,00-258,000 shooting days inland (PACEC 2006; PACEC 2014; Madden *et al*., 2025, Cameron *et al*. 2026,)). Despite the shortcomings of our approach, such “messy data” (Dobson *et al*. 2020), could be a useful resource in beginning to better understand harvest levels, especially in those countries without existing statutory harvest recording schemes. Those countries with established recording schemes are in a position to assess the suitability of this technique by comparing the proportions of species in their national bag recording schemes with those on photographs on social media. Our approach was not suitable for estimating the age demographics of harvested waterfowl, but may still be useful for sexing species and especially so for non-waterbird species with greater sexual dimorphism, including for example cervids and certain gamebirds.

## Data availability

The datasets generated during and/or analysed during the current study are at Datadryad (https://doi.org/10.5061/dryad.xwdbrv1qb)

