## Supplemental Table S1 for "Exploring sources of uncertainty in the estimate of waterfowl harvest in the United Kingdom"

Supplementary online information for **Exploring sources of uncertainty in the estimate of waterfowl harvest in the United Kingdom**

**Table S1**: Mean (± 95% Cis) harvest estimates for the United Kingdom of legally hunted wild duck species in 2016 when based on the proportions as seen on social media (Facebook), or the bag when assuming that for ducks those birds seen on social media are not linked to commercial hunting or rear and released game-farm mallards using estimates based on return rates from Aebischer (2024) (FB ≠ Released), or when release estimates are based on uncertainty in return rates from our own work (FB ≠ Released2) or in addition to FB ≠ Released2 including uncertainty in the total number of game-farm mallard released (FB ≠ Released3), or finally in addition to FB ≠ Released3 including uncertainty in the total number of wild duck harvested in the United Kingdom (FB ≠ Released3* total bag). These should be compared to those for the same species in the National Gamebag Census (Aebischer 2019). Data for Gadwall, Teal, Mallard and Wigeon are plotted in main manuscript (Figure 3).

| **Estimation Method** |  | **Gadwall** | **Teal** | **Mallard** | **Tufted** | **Wigeon** | **Shoveler** | **Pintail** | **Pochard** |
| --- | --- | --- | --- | --- | --- | --- | --- | --- | --- |
| Facebook | Mean | 55000 | 290000 | 670000 | 18000 | 89000 | 12000 | 6900 | 2600 |
|  | upperCI 97.5% | 74000 | 34000 | 760000 | 28000 | 120000 | 18000 | 11000 | 6100 |
|  | lowerCI 2.5% | 39000 | 23000 | 580000 | 9000 | 60000 | 7600 | 3700 | 330 |
| FB ≠ Released | Mean | 18000 | 97000 | 220000 | 5900 | 30000 | 4100 | 2300 | 890 |
|  | upperCI 97.5% | 26000 | 120000 | 280000 | 9800 | 43000 | 6200 | 3700 | 2100 |
|  | lowerCI 2.5% | 12000 | 71000 | 170000 | 2900 | 19000 | 1200 | 1200 | 120 |
| FB ≠ Released2 | Mean | 18000 | 96000 | 220000 | 5900 | 30000 | 4100 | 2300 | 890 |
|  | upperCI 97.5% | 27000 | 130000 | 290000 | 10000 | 44000 | 6200 | 3900 | 2100 |
|  | lowerCI 2.5% | 11000 | 65000 | 150000 | 2800 | 18000 | 2400 | 1100 | 100 |
| FB ≠ Released3 | Mean | 26000 | 140000 | 310000 | 8400 | 42000 | 5700 | 3200 | 1200 |
|  | upperCI 97.5% | 36000 | 170000 | 390000 | 14000 | 60000 | 8600 | 5300 | 2900 |
|  | lowerCI 2.5% | 17000 | 97000 | 240000 | 4100 | 27000 | 34000 | 1600 | 160 |
| FB ≠ Released3 * total bag | Mean | 26000 | 140000 | 310000 | 8400 | 42000 | 5700 | 3200 | 1200 |
|  | upperCI 97.5% | 48000 | 240000 | 550000 | 17000 | 78000 | 11000 | 6700 | 3400 |
|  | lowerCI 2.5% | 6300 | 33000 | 77000 | 1700 | 10000 | 1300 | 710 | 110 |
